# Beyond Blacklists: A Critical Assessment of Exclusion Set Generation Strategies and Alternative Approaches

**DOI:** 10.1101/2025.02.06.636968

**Authors:** Brydon P. G. Wall, Jonathan D. Ogata, My Nguyen, Amy L. Olex, Konstantinos V. Floros, Anthony C. Faber, Joseph L. McClay, J. Chuck Harrell, Mikhail G. Dozmorov

## Abstract

Short-read sequencing data can be affected by alignment artifacts in certain genomic regions. Removing reads overlapping these exclusion regions, previously known as Blacklists, help to potentially improve biological signal. Tools like the widely used Blacklist software facilitate this process, but their algorithmic details and parameter choices are not always clearly documented, affecting reproducibility and biological relevance. We examined the Blacklist software and found that pre-generated exclusion sets were difficult to reproduce due to variability in input data, aligner choice, and read length. We also identified and addressed a coding issue that led to over-annotation of high-signal regions. We further explored the use of “sponge” sequences—unassembled genomic regions such as satellite DNA, ribosomal DNA, and mitochondrial DNA—as an alternative approach. Aligning reads to a genome that includes sponge sequences reduced signal correlation in ChIP-seq data comparably to Blacklist-derived exclusion sets while preserving biological signal. Sponge-based alignment also had minimal impact on RNA-seq gene counts, suggesting broader applicability beyond chromatin profiling. These results highlight the limitations of fixed exclusion sets and suggest that sponge sequences offer a flexible, alignment-guided strategy for reducing artifacts and improving functional genomics analyses.

## Introduction

Alignment of short-read sequencing data to reference genome assemblies poses significant challenges due to the presence of low-complexity regions, centromeres, telomeres, satellite repeats, chromatin accessibility biases, and other artifacts [1]. These regions often result in abnormal read pileups caused by ambiguous alignments or biases introduced during library preparation steps, such as cell fixation or PCR amplification after adapter ligation [2,3]. Such artifacts have been observed across multiple species [4,5] and are particularly pronounced in chromatin-targeting sequencing technologies, including ChIP-seq [6,7,5], ChIP-exo [8], and CUT&RUN [9]. Removing these artifact signals has been shown to reduce noise and improve peak calling, signal normalization, and motif analysis accuracy [8,10,11,9,2]. Hence, defining strategies for removing such problematic signals is crucial for accurate genomic data analysis and interpretation.

Lists of exclusion regions (formerly known as blacklist regions), defined as genomic coordinates of problematic regions, have become a commonly accepted method for excluding artifact signals [12]. Several exclusion region sets have been developed for humans and model organisms, with an overview provided in [8]. The ENCODE project and others have compiled exclusion sets for various genome assemblies [13–15]. With technological advancements, such as long-read sequencing, sequencing data for newer organisms, and updated genome assemblies for existing organisms—such as the Telomere-to-Telomere (T2T) human genome assembly [16]— generating and updating these exclusion sets remains a priority. Notably, the commonly used liftOver tool for coordinate conversion has been deemed unsuitable for translating exclusion set coordinates across genome assemblies [13], highlighting the need for standardized exclusion set definitions and reproducible methods for generating them.

Methods for creating exclusion sets range from ad hoc approaches [17,5,9] to dedicated tools. Among these, the Blacklist software [13] is widely used. This tool employs BAM files from ChIP-seq input experiments (expected to have uniform non-specific sequencing coverage) and mappability files generated by Umap [18]. However, several key parameters of its algorithmic implementation are not fully documented, including the number of BAM files required, the read length and aligner specifications, and the k-mer length for selecting low-mappability files. The impact of these parameters on exclusion set generation remains unclear. Additionally, the software merges regions within a fixed distance of 20,000 bp—a parameter that appears excessive and is not easily adjustable. While these simplifications allow for straightforward use as a black-box tool, they raise concerns about the robustness and biological relevance of the resulting regions, limiting its applicability across sequencing technologies and reference genomes.

Alternative methods also have limitations. PeakPass employs a random forest model trained on hg19 ENCODE blacklist regions to predict excludable regions in hg38. It identifies assembly gaps and genome complexity as key predictors but lacks a user-friendly implementation. Similarly, Greenscreen [19] uses a traditional approach by processing ChIP-seq data through standard pipelines and calling peaks with MACS2 [20]. While Greenscreen claims 99% agreement with Blacklist-generated regions, its reliance on fixed parameters and a technically challenging Docker-based setup hampers its utility. The GreyListChIP R package [21], developed in 2015, defines excludable regions using a tiled genome and merges regions within a fixed distance. However, its unpublished algorithm and untested parameters further limit its reliability, particularly for less-studied organisms.

An alternative approach involves aligning sequencing data to reference genomes that include so-called “decoy” or “sponge” sequences. Such sequences typically consist of regions not included in the standard human reference genome. For example, the hs38d1 genome assembly (GCA_000786075.2) includes such sequences for the GRCh38/hg38 human genome assembly. A more comprehensive approach uses sponge sequences, which encompass unmapped and uncharacterized regions, satellite repeats, ribosomal sequences, and mitochondrial sequences. Incorporating these sponge sequences into the genome assembly during alignment has been shown to reduce signal in Blacklist exclusion regions and mitigate other alignment artifacts [1]. Despite its advantages, this approach is less widely adopted, with only 43 citations compared to 1,445 citations for the Blacklist manuscript (Google Scholar, February 2025).

In this study, we systematically benchmarked the performance of the Blacklist software [13] and related factors, such as aligner choice, to develop recommendations for exclusion set generation. Focusing on human and mouse genomes, we identified reproducibility and quality issues in existing exclusion sets. We provide a corrected, configurable version of the Blacklist software and propose improvements for defining exclusion regions. Rather than relying on fixed lists of exclusion regions, our results emphasize the importance of using “sponge” sequences at the alignment step. Our findings underscore the importance of transparent algorithms and adaptable methodologies to address the ongoing challenges in sequencing data analysis.

## Results

Filtering signal overlapping excludable (also known as Blacklist) regions is a standard practice in genomic data analysis. These regions were originally defined using input ChIP-seq data [8,13,22]. The simplicity of excludable regions (genomic coordinates in BED format) has enabled their application in analyzing genomic data generated by technologies based on assumptions other than ChIP-seq, such as ATAC-seq [23,24], chromatin conformation capture technologies [25], and their single-cell variants [26–28], as well as in pipelines and tools for genomic data analysis [29–31].

The ENCODE Blacklist exclusion set is arguably one of the most cited and frequently used exclusion sets for the human genome, and the Blacklist software implementation has been employed to generate exclusion sets for model organisms [13]. However, we have observed instances where signals expected to be associated with functionally relevant genes were missed, and such genes often overlapped with or were located in proximity to Blacklist exclusion regions. These observations prompted us to investigate the properties of the ENCODE Blacklist exclusion sets and to develop a set of recommendations for their optimal use.

### Challenges in reproducing pre-generated Blacklist exclusion sets

The Blacklist GitHub repository offers pre-generated exclusion sets (also referred to as lists) for human (hg19, hg38), mouse (mm10), Drosophila (dm3, dm6), and worm (cd10, ce11). We noted that version 1 of the hg38 exclusion set contained 38 regions, in contrast to 636 regions in version 2. In an effort to reproduce the hg38 “GitHub Blacklist,” we ran the software using the same set of 250 BAM files described in the original publication, generating what we refer to as the “Generated Blacklist.”

Our results indicated some differences compared to the GitHub version. Specifically, the “Generated Blacklist” contained more regions (1,273 vs. 636 in the GitHub version, Table 1), which were generally narrower (mean width approximately 213 Kbp vs. 357 Kbp in the GitHub version). The total genome coverage was also higher (approximately 271 Mbp vs. 227 Mbp in the GitHub version, Figure 1A), with similar telomere, centromere, and short-arm coverage (referred to hereafter as gap coverage), except for the short arms, which were primarily covered by our generated list (Figure 1B). Most regions from the “GitHub Blacklist” overlapped with the “Generated Blacklist” (79.4%, covering 97.1% of the same bases, Figure 1B). However, both lists showed notable differences from the manually curated “Kundaje Unified” list, which is recommended as a manually curated gold standard [12]. Only 18.1% of regions from the “GitHub Blacklist” overlapped with regions from the “Kundaje Unified” list, with overlap by width similarly limited (28.8%, Figure 1C).

**Table 1.**
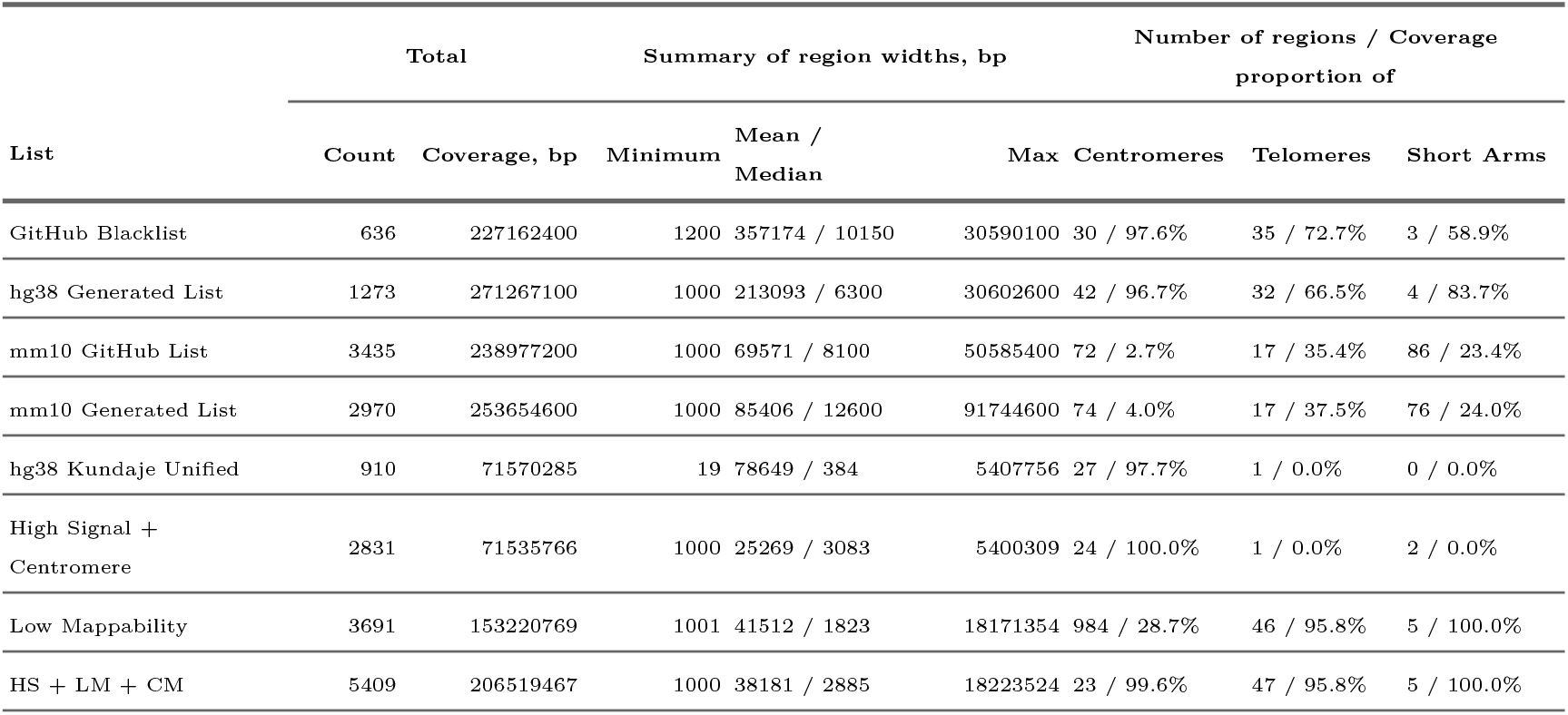
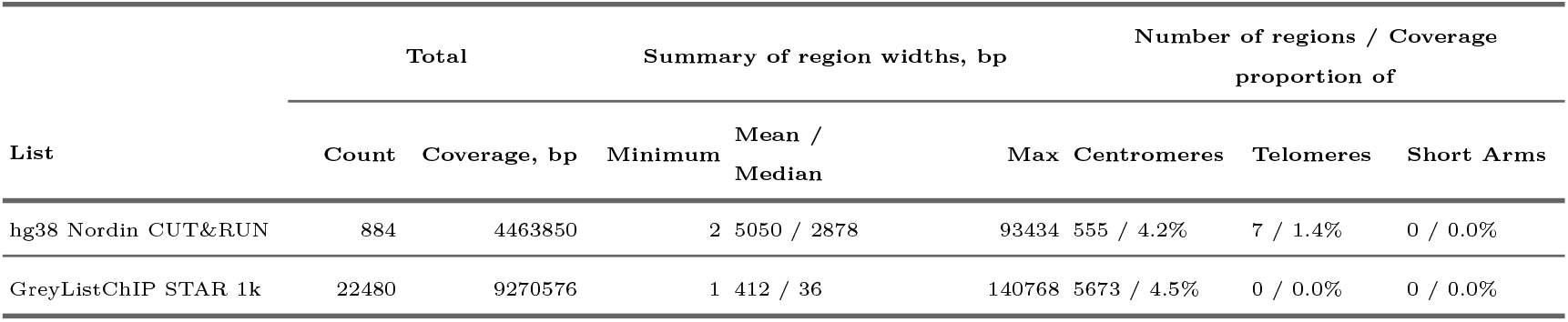
Characteristics of the hg38 exclusion sets. Total number of regions, average region width, number of regions overlapping gaps, and proportion of gap coverage.

**Figure 1.**
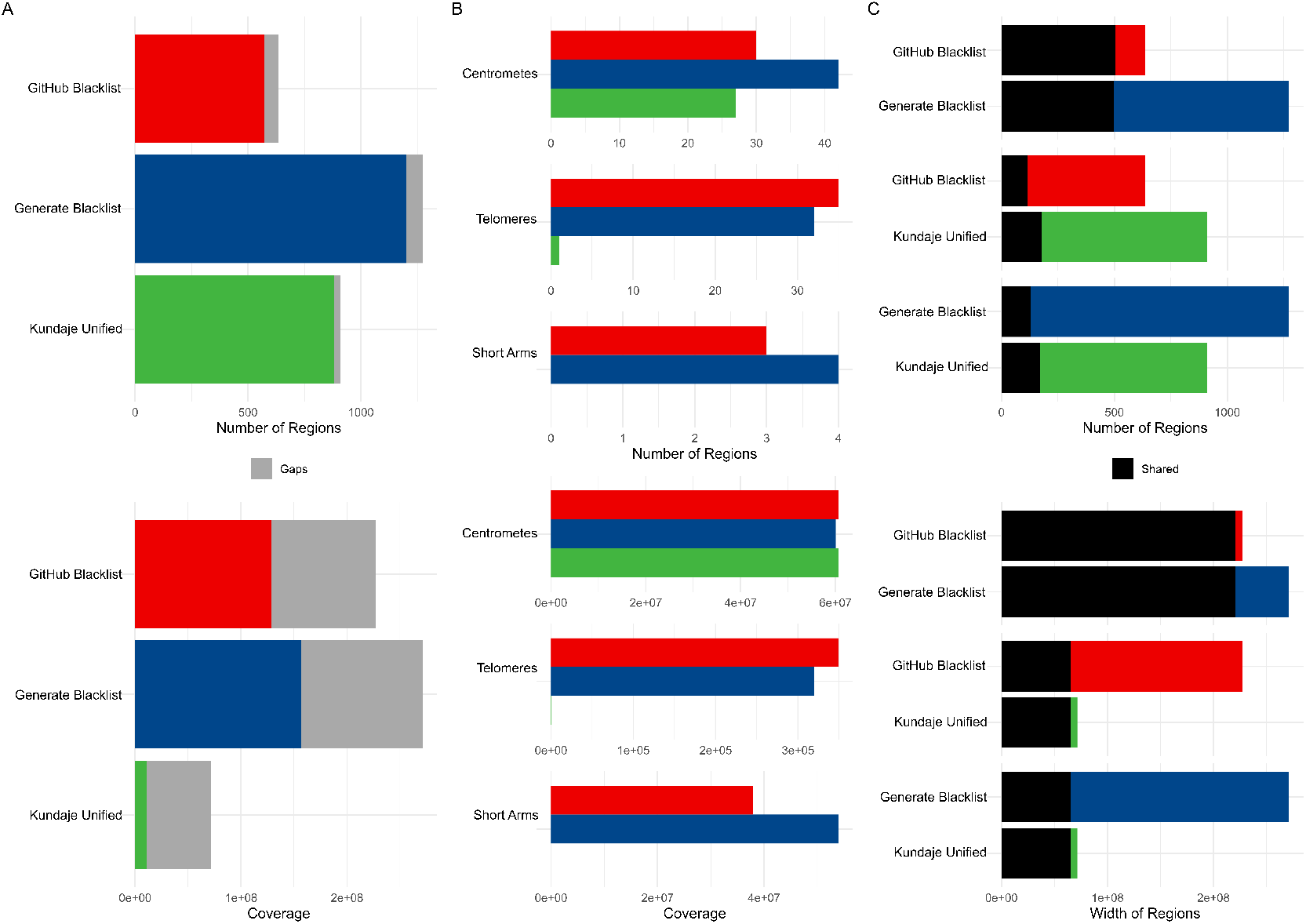
Differences between the GitHub version of the hg38 exclusion set, our hg38 exclusion set generated with the Blacklist software, and the reference Kundaje Unified set. A) Count and coverage differences; B) Gap coverage differences; (C) Pairwise overlaps of region counts and genome coverage between sets.

For the mm10 list, our results showed fewer regions compared to the GitHub version (2,970 vs. 3,435), but the regions were wider on average (49,076 bp vs. 41,235 bp) and covered more bases (approximately 253 Mbp vs. 238 Mbp in the GitHub version) (Supplementary Figure S1A, Additional File 1, Supplementary Table S1, Additional File 2). The overall gap coverage between the two mouse Blacklists was comparable (Supplementary Figure S1A, Additional File 1). While approximately 30% of regions were unique to each list, the shared regions accounted for over 80% of common base coverage (Supplementary Figure S1C, Additional File 1).

To support reproducibility, we have made our analysis scripts available (Code Availability). These findings highlight some variability in exclusion sets generated by the Blacklist software and suggest the value of cross-comparison with curated references like the “Kundaje Unified” list.

To explore the differences between the exclusion sets provided on GitHub and those we generated, we examined the properties of the 1,255 BAM files used to create the hg38 “GitHub Blacklist.” Our analysis indicated that a significant portion of the BAM files (37.6%) originated from 36 bp-long reads, followed by 101 bp-long reads (21.8%). However, read length varied considerably, with files generated from reads of 28 bp, 50 bp, 76 bp, and other lengths (Supplementary Figure S2A, Additional File 3, Supplementary Table S2, Additional File 4). Additionally, 365 out of 1,255 files (29.1%) were derived from paired-end experiments and aligned using bwa sampe v.0.7.10, while single-end files were aligned with bwa samse. The average number of mapped reads also differed, with paired-end files having approximately 89 million mapped reads, compared to 38 million for single-end files (Supplementary Figure S2B, Additional File 3).

We observed further variation in the dataset, including 256 files that combined multiple (pooled) FASTQ files and 60 that used cropped FASTQ files, where the cropping parameters were not specified. Of the total BAM files, 135 were restricted (FASTQ sequences unavailable), 52 contained duplicated FASTQ files, and 4 were both restricted and duplicated (Supplementary Table S2, Additional File 4). These BAM files were merged into 250 donor-specific files in the original publication [13], with the number of BAM files per donor ranging from 1 to 149 (average of 5) (Supplementary Figure S2C, Additional File 3). We also noted variability in donor-specific BAM files, which differed in read length, mixed single- and paired-end sequencing, and were derived from various cell and tissue types (Supplementary Figure S2D, Additional File 3, Supplementary Table S2, Additional File 4). To assess whether merging files affected exclusion set generation, we compared exclusion sets generated from 274 101 bp paired-end BAM files (spanning 21 donors) to those produced from a single BAM file containing all merged 101 bp data. The differences between the resulting exclusion sets were minimal (Supplementary Figure S3, Additional File 5).

These observations suggest that the diversity in input data characteristics, along with differences in sequencing and alignment protocols, may influence the reproducibility and consistency of exclusion sets generated by the Blacklist software. By highlighting these factors, we hope to encourage further consideration of input data variability in exclusion set generation.

### Manually defined High Signal and Low Mappability regions are different from other exclusion sets

To evaluate the reproducibility of exclusion sets in the absence of a definitive gold standard for excludable regions, we chose the manually curated “Kundaje Unified” list as a reference for known hg38 exclusion regions. This list includes a larger number of regions compared to the GitHub-provided exclusion set (910 regions in the “Kundaje Unified” list vs. 636 in the GitHub version, Figure 2A). Of these, 179 regions overlapped with the GitHub list (Figure 2B, Supplementary Figure S4B, Additional File 6). The “Kundaje Unified” regions were generally narrower (mean width of 78.6 Kbp compared to 357.2 Kbp in the GitHub list, Figure 2C) and covered less of the genome (71.6 Mbp for the “Kundaje Unified” list vs. 227.2 Mbp for the GitHub list, Table 1). While the “Kundaje Unified” list provided near-complete coverage of centromeres (97.7%), it did not extend to telomeres or short arms (0.0%, Supplementary Figure S4C, Additional File 6). In contrast, the GitHub list covered 97.6% of centromeres, 72.7% of telomeres, and 58.9% of short arms (Supplementary Figure S4C, Additional File 6). For our benchmarking, we used the “Kundaje Unified” list as the primary reference, recognizing its role as a curated and widely referenced set of exclusion regions.

**Figure 2.**
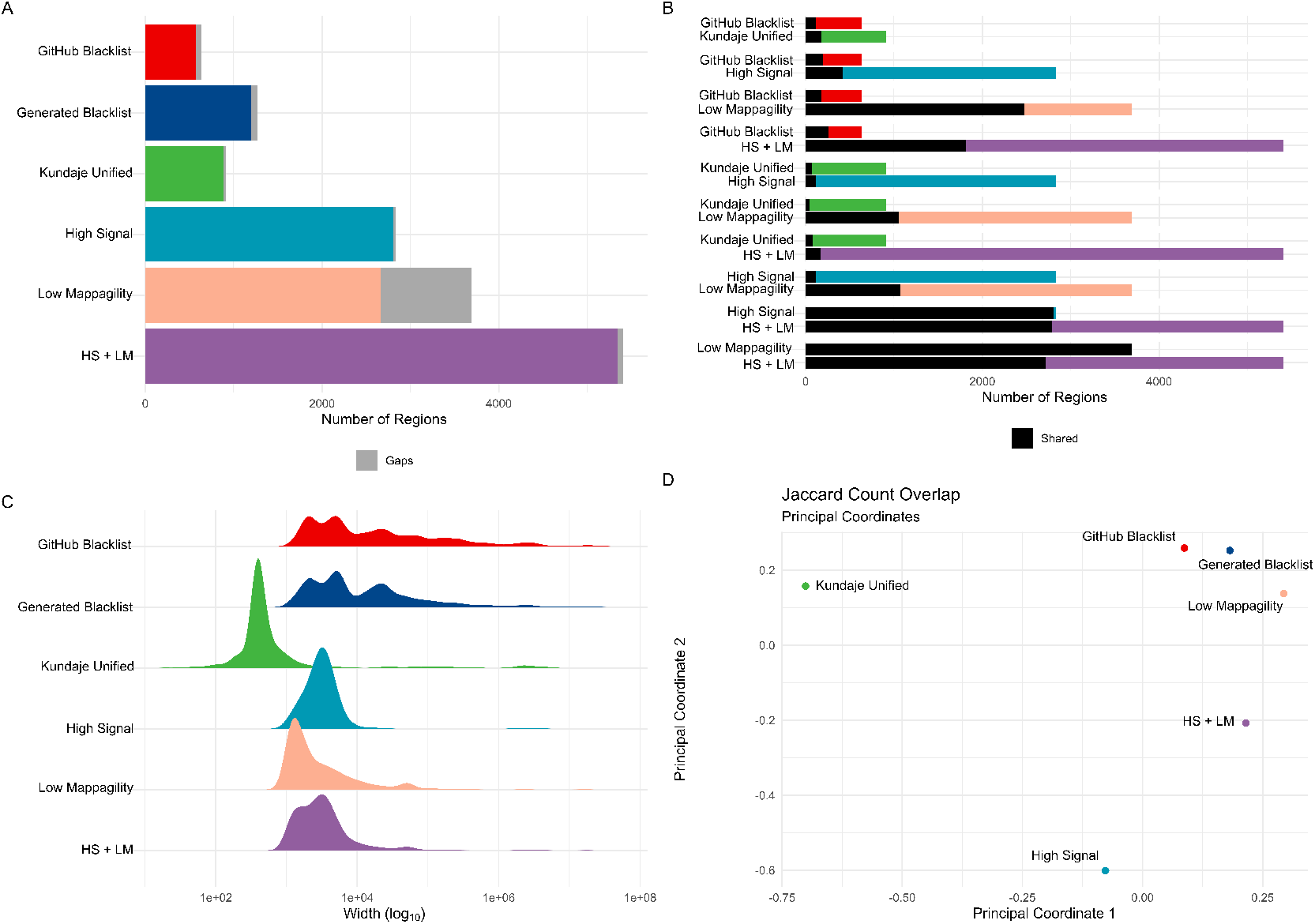
Differences between the GitHub version of the hg38 exclusion set and manually defined gold-standard exclusion sets. (A) Differences in count. (B) Pairwise overlaps of region counts between the GitHub exclusion set and manually defined gold-standard exclusion sets. (C) Differences in width distribution. (D) Multidimensional scaling plot of Jaccard count similarity among the GitHub exclusion set and manually defined gold standards.

High Signal (HS) regions are another well-characterized category of exclusion regions [24]. Using 274 101 bp paired-end BAM files, we identified “High Signal” regions by applying MACS3 to call peaks with fold changes exceeding the 99th percentile. We merged regions within 1,000 bp and excluded those smaller than 1,000 bp (Methods). Our analysis showed that the “High Signal” regions had limited coverage of centromeres compared to the “Kundaje Unified” list (2.1% vs. 97.7%). To address this, we incorporated centromeric regions into our “High Signal” list, producing a final set of 2,657 regions (Figure 2A). This adjusted list covered a comparable portion of the genome and gap regions as the “Kundaje Unified” list (71.0 Mbp vs. 71.6 Mbp, Supplementary Figures S4A and S4B, Additional File 6). The “High Signal” regions were generally narrower than those in the “Kundaje Unified” list and the GitHub Blacklist (mean width of 26.7 Kbp, Figure 2B). While 422 “High Signal” regions overlapped with the GitHub Blacklist, accounting for 27.3% of its total coverage, the overlap with the “Kundaje Unified” list was smaller in number (113 regions) but encompassed a significant portion of its base coverage (85.2%, Figure S4C). This “High Signal” list offers a conservative representation of high-signal regions derived from input ChIP-seq data.

Low Mappability (LM) regions represent another category considered by the Blacklist software. Following an approach similar to that used for “High Signal” regions, we defined “Low Mappability” regions as those falling below the 1st percentile of the mappability range. Regions within 1,000 bp were merged, and regions smaller than 1,000 bp were excluded (Methods). Despite applying this conservative strategy, we identified a substantial number of “Low Mappability” regions (12,455, Figure 2A), which tended to be wide (mean width of 41.5 Kbp, Figure 2B). These regions covered a significant portion of the genome (153.2 Mbp, Figure S4A), including centromeres (28.7% coverage), telomeres (95.8%), and short arms (100%, Supplementary Figure S4C, Additional File 6). The “Low Mappability” regions overlapped with 228 regions from the GitHub Blacklist, accounting for 51.3% of its coverage (Figure 2C). They also intersected with 45 regions from the “Kundaje Unified” list, covering 26.7% of its bases (Figure 2C). This “Low Mappability” list reflects a conservative set of regions that are challenging to map, and it appears to differ from both the GitHub Blacklist and the “Kundaje Unified” lists.

To align with the characteristics of the Blacklist-generated list, which includes both high signal and low mappability regions, we combined the “High Signal” (with centromeres) and “Low Mappability” lists, referred to as “HS + LM”. This resulted in 5,409 regions with a mean width of 38.1 Kbp, covering 206.5 Mbp of the genome and nearly complete gap coverage (Figure 2A–B, Supplementary Figure S4A–B, Additional File 6, Table 1). Although 2,078 (38.4%) of the “HS + LM” regions overlapped with the GitHub Blacklist, they accounted for 70.6% of its total coverage. Similarly, while only 173 (3.2%) regions overlapped with the “Kundaje Unified” list, they covered 87.0% of it, largely driven by centromeric regions (Figure 2C, Supplementary Figure S4C, Additional File 6). The “HS + LM” list, created as a single reference for addressing genomic regions prone to mapping artifacts, highlights distinctions from both the GitHub Blacklist and the “Kundaje Unified” list.

To explore similarities between exclusion lists, we visualized Jaccard overlap indexes among them using multidimensional scaling (Methods). The “GitHub Blacklist” and our “Generated Blacklist” showed similarity, while the “Kundaje Unified,” “High Signal,” “Low Mappability,” and “HS + LM” lists appeared more distinct (Figure 2D). To account for differences in list size, region number, and genome coverage, we applied the Forbes width overlap coefficient, known for its robustness in such contexts [32]. This analysis indicated that the “Kundaje Unified” and “High Signal” lists share some similarities and align more closely with the “GitHub Blacklist,” whereas the “Low Mappability” and “HS + LM” lists diverge (Supplementary Figure S4D, Additional File 6). These findings highlight notable heterogeneity among exclusion lists, suggesting that no single reference set universally defines problematic regions. Moving forward, we benchmark the Blacklist software using the “Kundaje Unified” list alongside the “High Signal,” “Low Mappability,” and “HS + LM” lists to provide a broader evaluation of performance.

### Excessive High Signal annotation of Blacklist-generated regions

Exclusion sets produced by the Blacklist software include annotations indicating whether regions are classified as “High Signal” or “Low Mappability,” with no overlap between categories. We observed that “High Signal” annotations were predominant across all organisms’ lists (83.75 *±* 9.54%), with the hg38 “GitHub Blacklist” showing a particularly strong bias (93.40%, Supplementary Table S2, Additional File 4). To investigate this pattern, we examined the Blacklist software’s algorithm and original C code (Supplementary Note, Additional File 7, Supplementary Figure S5, Additional File 8). Our analysis revealed that the software defaulted to “High Signal” annotations in cases of ambiguity, as it could not output combined classifications. We addressed this by modifying the code to allow for a new annotation, “High Signal, Low Mappability,” and have made the updated version available (Code Availability).

The modified code produced consistent results (Supplementary Table S1, Additional File 2) and, for the hg38 genome assembly, identified the same 65 “Low Mappability” regions spanning 7.4 Mbp. Of the 1,208 “High Signal” regions (covering 263.8 Mbp), 183 were reclassified as “High Signal, Low Mappability.” Although these reclassified regions were few, they accounted for 219.4 Mbp, representing 80.88% of the total genome coverage in the Blacklist-generated hg38 exclusion set. Similar trends were observed for the mm10 assembly (Supplementary Table S1, Additional File 2). These findings underscore the importance of recognizing multi-annotation regions in exclusion set generation and highlight the potential for annotation biases to influence downstream analyses.

### The choice of aligner and read length affects exclusion set generation

We hypothesized that the choice of aligner could influence the resulting exclusion sets. To assess this effect, we used STAR, bwa-mem2, and bowtie2 aligners (Methods), along with the originally used bwa samse, to realign 381 36 bp single-end files and generate exclusion sets using the Blacklist software. The 36 bp read length was selected to minimize variability from read length differences and to align with the internal k-mer setting of 36 (Supplementary Note, Additional File 7). We compared these exclusion sets with the “GitHub Blacklist” and “Generated Blacklist,” which were derived from all files aligned using bwa samse, and included the “Kundaje Unified” list as a reference. Realigned sets generally contained a larger number of regions. Notably, the STAR-realigned set had a region count closest to the “GitHub Blacklist” (749 vs. 636), while the bwa-mem2-realigned set had the highest number of regions (1,174, Supplementary Table S3, Additional File 9). The STAR-realigned set also exhibited a width distribution most similar to the “GitHub Blacklist” (mean width 381 kbp vs. 357 kbp), whereas sets from other aligners had narrower distributions (Figure 3A). Realigned sets covered a broader portion of the genome, including gap regions (Figure 3B). They overlapped approximately 60–80% of the regions in the “GitHub Blacklist” and covered up to 99% of its width (Figure 3C). Jaccard count multi-dimensional scaling and hierarchical clustering demonstrated that the STAR-aligned set closely resembled the “GitHub Blacklist,” while sets from other aligners were more similar to each other and distinct from both the “GitHub Blacklist” and the “Kundaje Unified” list (Supplementary Figure S6A, B, Additional File 10). Forbes width overlap plots also highlighted differences between the realigned sets and the reference lists (Supplementary Figure S6C, D, Additional File 10). These findings underscore the potential influence of aligner choice on exclusion set generation and highlight the importance of considering alignment tools in such analyses.

**Figure 3.**
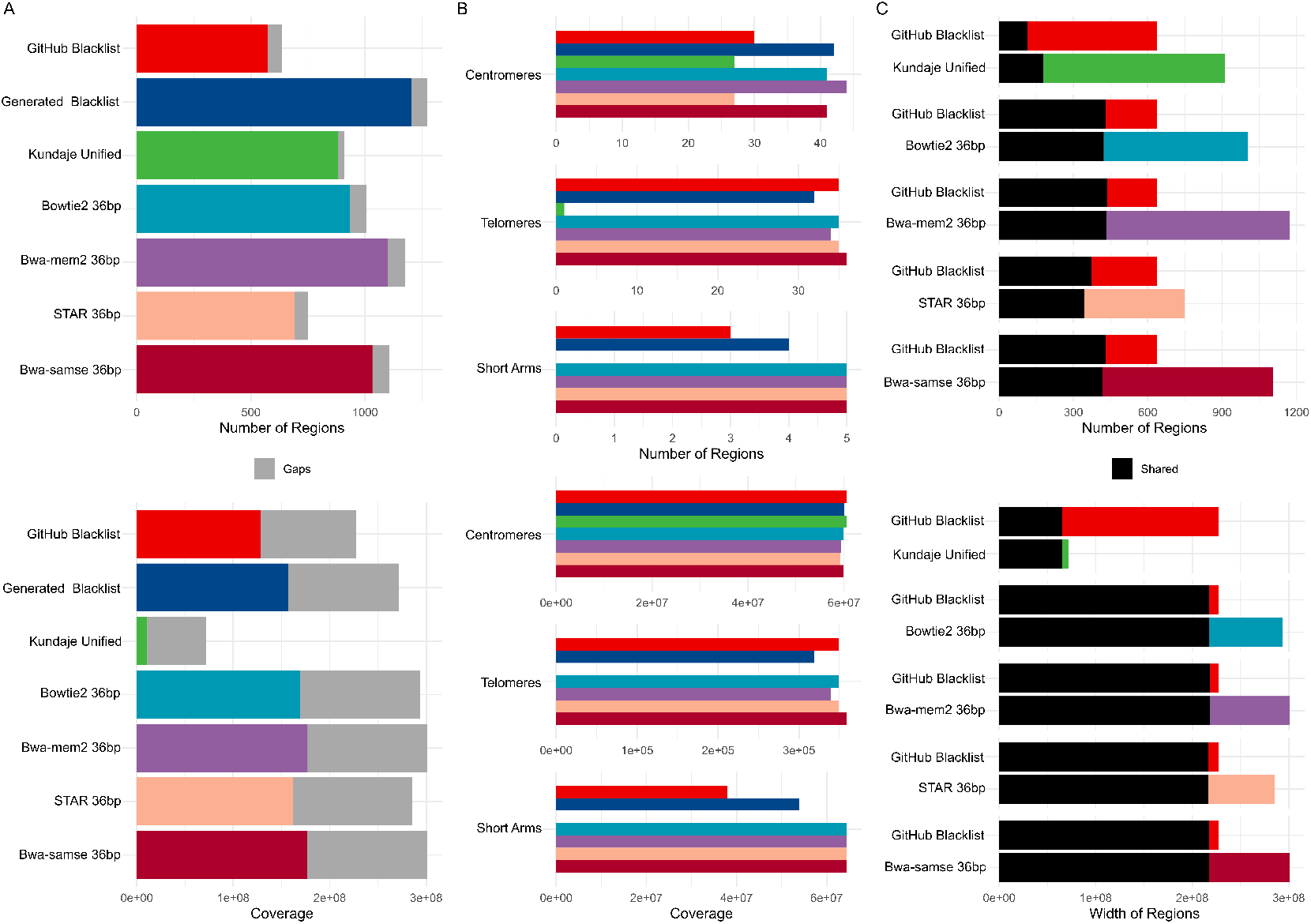
Differences between the hg38 exclusion sets, the reference Kundaje Unified set, and sets generated from 36bp single-end reads realigned with different aligners. A) Count and coverage differences; B) Gap coverage differences; C) Pairwise overlaps of region counts and genome coverage between sets.

We further hypothesized that read length could influence the resulting exclusion sets. To test this, we realigned 274 101 bp paired-end files using the same set of aligners and compared the resulting exclusion sets. The use of 101 bp files was associated with a larger number of regions that were generally narrower and covered a broader portion of the genome (Supplementary Figure S7A, Additional File 11, Supplementary Table S3, Additional File 9). Examining the impact of aligners, bwa-mem2 consistently identified a larger number of regions, which covered a greater portion of the genome but tended to be narrower. Conversely, STAR identified fewer regions that covered a smaller portion of the genome but were generally wider (Supplementary Table S3, Additional File 9). Sets generated with STAR were distinct from those generated using bwa-mem2 and bowtie2. However, the segregation of sets was primarily influenced by read length, with 36 bp STAR-generated exclusion sets showing the highest similarity to the “GitHub Blacklist” (Supplementary Figure S7B, Additional File 11). These findings suggest that the Blacklist software is sensitive to the read length of input BAM files, emphasizing the importance of considering read length in exclusion set analyses.

### Parameter selection for exclusion set calling

Recognizing the sensitivity of the Blacklist software to input settings, we systematically evaluated the effects of three parameters on the resulting exclusion sets. These parameters included the number of files (ranging from 10 to 300) to determine the optimal number needed for exclusion set generation, the “bridge” parameter (tested at 1,000, 10,000, and the default 20,000) to assess the optimal merging distance for nearby exclusion regions, and the “k-mer” parameter (default 36, 50, and 100) to evaluate the ideal k-mer size. For this analysis, we used the “Kundaje Unified,” “High Signal,” “Low Mappability,” and “HS + LM” sets as references, along with the 381 36 bp single-end STAR-aligned BAM files, which produced exclusion sets most similar to the hg38 Blacklist (Supplementary Figure S7B, D, Additional File 11). We measured the effects of these parameters on the number of regions, total coverage, and mean/median width.

Increasing the number of files and the “bridge” parameter generally resulted in fewer but wider regions that covered a larger portion of the genome. In contrast, increasing the k-mer parameter led to a greater number of narrower regions that also covered a larger genomic portion (Supplementary Figure S8, Additional File 12, Supplementary Table S4, Additional File 13). Jaccard count overlap and Forbes width similarity analyses revealed distinct parameter effects. The k-mer parameter had the most significant impact on Jaccard count overlap similarity (Supplementary Figure S9C, Additional File 14), followed by the number of BAM files (Supplementary Figure S9A, Additional File 14). The “bridge” parameter had the greatest influence on Forbes width similarity (Supplementary Figure S9F, Additional File 14), followed by the k-mer parameter (Supplementary Figure S9G, Additional File 14). Interestingly, none of the parameter combinations produced exclusion sets closely resembling the “Kundaje Unified” or the “High Signal,” “Low Mappability,” and “HS + LM” reference lists. However, exclusion sets generated using only 10 BAM files were unexpectedly the closest to the GitHub list by Jaccard count overlap similarity (Supplementary Figure S9D, H, Additional File 14). These findings underscore the sensitivity of the Blacklist software to input parameters and highlight its limitations in replicating classical definitions of excludable regions, such as high-signal or low-mappability peaks.

### Ribosomal genes are most affected by exclusion regions

To assess the impact of different exclusion sets on known transcripts, we quantified the extent to which these sets overlapped with transcripts (Supplementary Table S5, Additional File 15). Our analysis included two newer exclusion sets: the “Nordin CUT&RUN” list [33] and the “GreyListChIP” list, generated using the GreyListChIP R package with a 1,000 bp merging setting. These two lists exhibited similarity to each other but were distinct from other sets, except for the “Nordin CUT&RUN” list, which showed some similarity to the “GitHub Blacklist” (Supplementary Figure S10, Additional File 16). We observed that the “GreyListChIP” list and the “High Signal” list affected the largest number of protein-coding genes, whereas the “Generated Blacklist” covered the greatest portion of gene bases (Figure 4A). In contrast, the “Kundaje Unified” list had the smallest impact on the number and coverage of protein-coding genes, likely reflecting its manual curation, which intentionally avoids regions overlapping transcripts. Similarly, the “Nordin CUT&RUN” list showed a low impact on both the number and coverage of protein-coding genes. Comparable trends were observed for long noncoding RNAs and other transcript types (Figure 4A).

**Figure 4.**
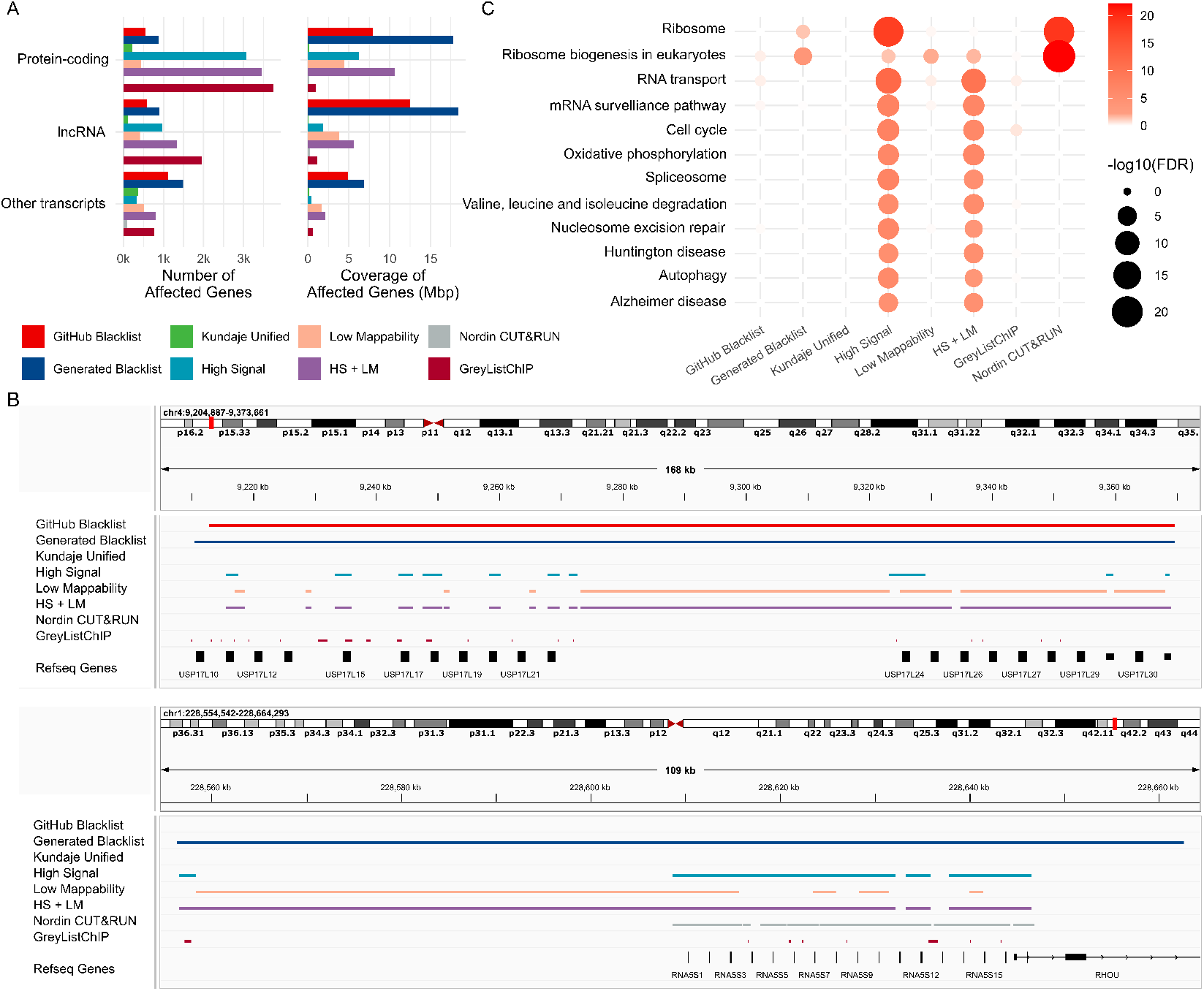
Biological characterization of genes affected by exclusion regions. A) Count and coverage of protein-coding, long noncoding, and other transcripts affected by exclusion sets; B) Representative comparison of exclusion set coverage over a cluster of ubiquitin-specific peptidase 17-like family member genes; C) KEGG pathways enriched in genes overlapped by exclusion sets.

We visualized a representative region of ubiquitin-specific peptidase 17-like family member genes to illustrate the large size of the Blacklist-generated regions, the “Low Mappability” regions that harbor protein-coding genes, the tendency of “High Signal” regions to overlap genes, and the conservative approach of the “Kundaje Unified” and the “Nordin CUT&RUN” lists, which avoid overlapping genes (Figure 4B, upper panel). These results indicate that the Blacklist-generated list, “High Signal,” “Low Mappability,” and the combined lists may remove peaks overlapping gene regions.

To determine whether specific gene sets or pathways are affected by exclusion regions, we performed KEGG enrichment analysis (Supplementary Table S5, Additional File 15). Pathways related to “Ribosome” and “Ribosome biogenesis in eukaryotes” were enriched in genes affected by the “Nordin CUT&RUN” list, the “High Signal” regions, the combined “HS + LM” regions, and our “Generated Blacklist” (Figure 4C). Genes from these pathways were also affected by the “Low Mappability” and “GitHub Blacklist” regions, though the observed enrichments were not statistically significant. This result was expected, as ribosomal RNA gene (rDNA) repeats are a well-characterized source of contamination [34–36], and ribosomal RNA (rRNA) constitutes approximately 80–90% of total cellular RNA [37]. However, the “GitHub Blacklist,” “Kundaje Unified,” and “GreyListChIP” lists did not show any significant enrichment.

We visualized a cluster of ribosomal genes affected by the “Nordin CUT&RUN” list (Figure 4B, lower panel), highlighting a key distinction between the “GitHub Blacklist” and the “Generated Blacklist.” The latter covered ribosomal genes, whereas the former did not, suggesting that the “GitHub Blacklist” may have been filtered to minimize overlap with gene regions. These findings underscore the unique characteristics of the “Nordin CUT&RUN” list, which, despite affecting fewer genes overall, appears to exhibit exclusive enrichment in ribosomal genes.

We examined the impact of exclusion sets on cancer-associated genes using oncoEnrichR analysis [38]. The analysis identified five oncogenes and tumor suppressors (collectively referred to as cancer drivers) with moderate to very strong evidence of being affected by the “GitHub Blacklist,” with KDM5A (lysine demethylase 5A) and MLH1 (mutL homolog 1) being the most notable oncogene and tumor suppressor, respectively (Supplementary Table S6, Additional File 17). The “Generated Blacklist” impacted 15 cancer drivers, including the NOTCH2 oncogene and tumor suppressor, with very strong confidence. The “High Signal” list affected 24 cancer drivers, encompassing key genes such as PIK3CA, MTOR, RAF1, and JUN. The “Low Mappability” list impacted six cancer drivers, including NOTCH2 and SSX2 (SSX family member 2). The combined “HS + LM” list affected the highest number of cancer drivers (32), while the “GreyListChIP” list impacted 17, including NOTCH2 and TP53. In contrast, the “Kundaje Unified” and “Nordin CUT&RUN” lists did not overlap with cancer driver genes. These observations suggest that the “High Signal” and “Low Mappability” regions should not be excluded and position the “Kundaje Unified” list as a viable option for analyses aiming to minimize the impact on functional pathways and oncogenes.

### Using “sponge” sequences decreases artificial correlation in ChIP-seq data comparable to the Blacklist-generated exclusion sets

The original Blacklist publication suggested that removing reads overlapping exclusion regions would enhance the biological interpretability of ChIP-seq data by reducing signal correlation among ChIP-seq profiles. The authors hypothesized that read pileups in exclusion regions lead to spurious correlations and that removing them would allow for better biological interpretability. In this study, we evaluated this hypothesis by correlating ChIP-seq transcription factor signals for the GM12878 cell line before and after removing reads overlapping exclusion regions. Notably, removing reads overlapping the “High Signal,” “Low Mappability,” and “Kundaje Unified” lists resulted in the smallest decrease in correlations (Figure 5A). We confirmed that removing reads overlapping the “Generated Blacklist” and the “GitHub Blacklist” lists resulted in the largest decrease in signal correlation for the REST, SREBF2, and CTCF transcription factors (Figure 5B).

**Figure 5.**
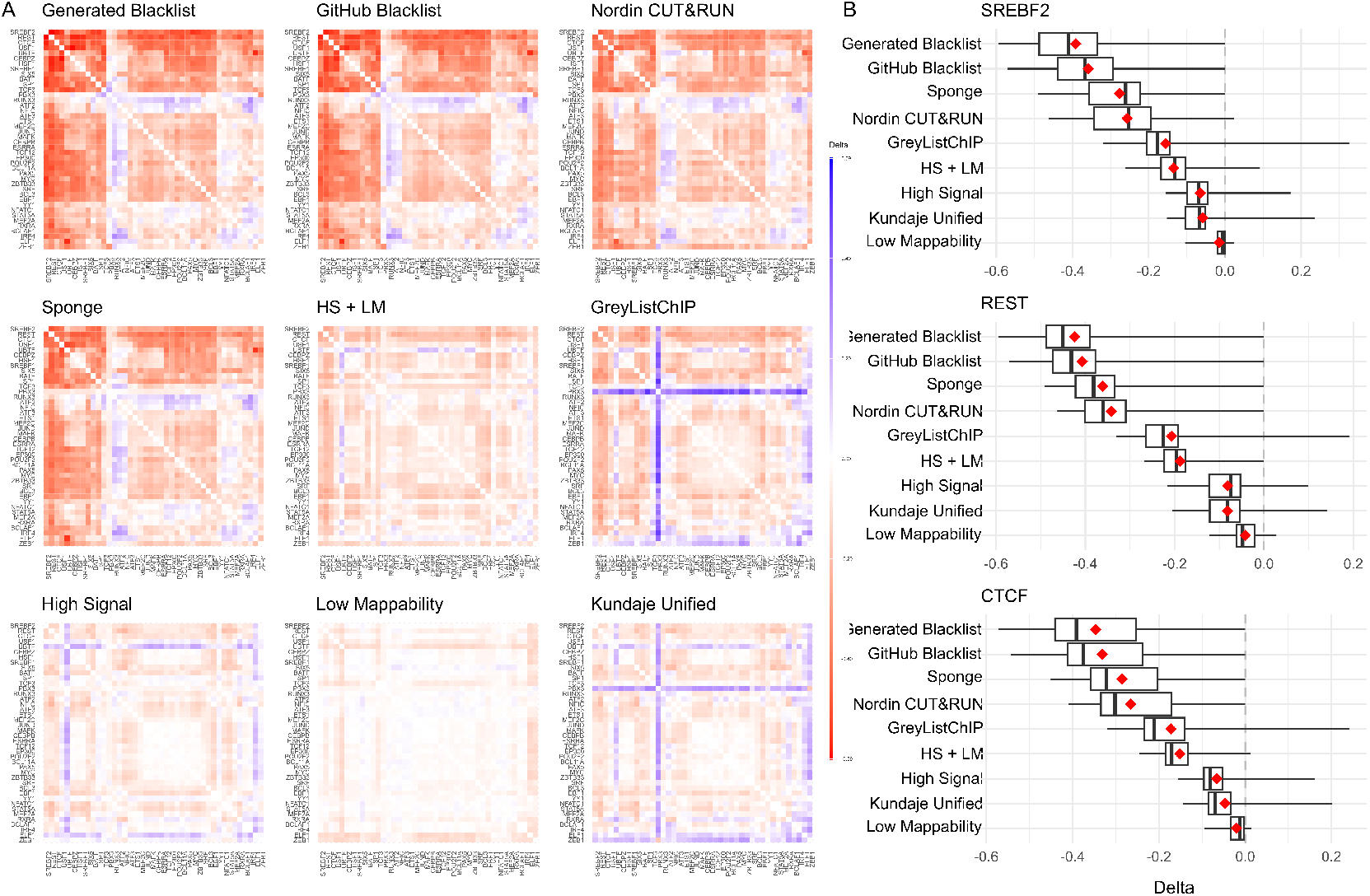
Changes in ChIP-seq signal correlation with and without reads overlapping exclusion sets or aligned to the sponge. Data for the Gm12878 cell line is shown. A) Heatmaps of correlation differences for each exclusion set, sorted by mean (red/blue gradient corresponds to decreases/increases in correlations, respectively); B) Correlation difference distributions for the top three most affected transcription factors.

Aligning sequencing reads to a genome that includes so-called “sponge” sequences represents another strategy to reduce artifact signals. These sequences include satellite DNA, ribosomal DNA, mitochondrial sequences, etc., and constitute roughly 8% of the human genome omitted from the reference assembly [1]. We hypothesized that including these “sponge” sequences at the alignment step would reduce read pileups and signal correlation compared to reads aligned to the assembled chromosomes only. We found that this was indeed the case, with the overall “sponge”-aligned signal correlation reduction being second only to the “Generated Blacklist” and “GitHub Blacklist” lists (Figure 5). With minor exceptions, these observations were consistent for other transcription factors (Supplementary Figure S11, Additional File 18).

We investigated the alignment characteristics of reads aligned to the “sponge” sequences and found that 48.78% of 36 bp-long reads were aligned to them (Supplementary Figure S12A, Additional File 19, Supplementary Table S7, Additional File 20). In contrast, only 6.21% of 101 bp-long reads were aligned to the “sponge” sequences, suggesting that the higher information content in longer reads improves alignment to the autosomal reference. We also investigated the number and proportion of reads derived from excludable regions that overlapped the “sponge” sequences and found that the “Nordin CUT&RUN” and “GreyListChIP” lists had the largest proportion of such reads, followed by the “Kundaje Unified” list (80.38%, 74.30%, and 45.66%, respectively; Supplementary Table S7, Additional File 20). In summary, these results position the “sponge” alignment as a viable alternative to fixed region-based read filtering.

### “Sponge” sequences preserve biological signal in omics data

We hypothesized that using the “sponge” sequences would minimally affect or improve biological signals in omics data. We utilized two versions of the hg38 genome assembly: one that included only autosomal sequences, and another, referred to as “full plus sponge,” that additionally included unplaced contigs and the “sponge” sequences. To investigate whether the telomere-to-telomere human genome assembly could substitute for the effect of “sponge” sequences and contigs, we further utilized the T2T-CHM13 genome assembly [16].

We analyzed RNA-seq data from patient-derived xenograft (PDX) models of breast cancer and found that the total number of reads overlapping gene regions remained nearly unchanged (Supplementary Figure S13A, Additional File 21; p-value > 0.3, pairwise Wilcoxon test). PCA analysis also demonstrated that the alignment strategy preserved biological signal and did not introduce batch effect-like variability (Supplementary Figure S13B, Additional File 21). We performed differential gene expression analysis and found that inclusion of contigs and “sponge” sequences in the hg38 genome assembly resulted in significant downregulation of six genes (three noncoding and three ribosomal). Comparison of data aligned to the T2T-CHM13 assembly versus the hg38 assemblies resulted in 1,392/1,377 upregulated and 495/503 downregulated genes for hg38 autosomes vs. T2T and hg38 full plus sponge vs. T2T, respectively. These gene sets were highly overlapping (Supplementary Figure S13C, Additional File 21), and the majority of them (56%) were ncRNAs, while only 34% were protein-coding (Supplementary Table S8, Additional File 22). To understand their functional significance, we performed KEGG pathway enrichment analysis. We found clusters of histone genes associated with the immune system upregulated in the T2T-CHM13–aligned data (Supplementary Table S8, Additional File 22). This is expected, as histone gene regions are highly repetitive, contain multiple highly similar copies, and were difficult to resolve in the hg38 genome assembly [16]. Among downregulated genes, we observed members of the Speedy/RINGO cell cycle regulator family, which encode cyclin-like proteins that bind and activate cyclin-dependent kinases (CDKs) and play roles in cell cycle and growth (Supplementary Table S8, Additional File 22). This is also expected, as the SPDYE gene family consists of multiple tandemly arrayed paralogs resolved in the T2T genome assembly, making them a high-copy, highly similar cluster that was hard to resolve in hg38 [39]. These results highlight the potential for T2T-CHM13 alignment to provide richer biological insights, but also demonstrate the overall minimal impact of genome assembly choice and “sponge” sequences on the transcriptome.

ATAC-seq represents another data type that may benefit from alignment with “sponge” sequences. We compared the number of peaks differentially accessible between treated and untreated conditions (see Methods) and found that including alternative contigs and “sponge” sequences reduced the number of peaks by approximately 3.7% (49,496/47,583 peaks in hg38 autosomal/full plus sponge alignment settings). Alignment to the T2T-CHM13 genome assembly resulted in the largest number of peaks (49,538; Supplementary Table S9, Additional File 23). This is expected, as the T2T-CHM13 genome assembly should include previously unplaced contigs and/or sponge sequences. Peaks detected under each alignment strategy had similar widths (median 248 bp, t-test p-value > 0.3). We annotated peaks with genes and found the majority overlapped between the hg38 autosomal and full plus sponge alignments (Supplementary Figure S13D, Additional File 21). Alignment to the T2T-CHM13 genome assembly resulted in the largest number of unique genes (6,360), 305 of which were protein-coding. The hg38 autosomal-only alignment resulted in peaks associated with 195 unique genes (53 protein-coding). By contrast, the hg38 full plus sponge alignment resulted in 54 unique genes (28 protein-coding; Supplementary Figure S13D, Additional File 21). To assess the functional importance of these unique genes, we performed KEGG pathway enrichment analysis. No significant enrichments were found in any of the unique gene sets, suggesting that different alignment strategies do not affect specific functional categories. However, closer inspection of the unique protein-coding genes in the hg38 full plus sponge alignment revealed several clinically relevant genes, such as FOS (oncogene, AP-1 complex), BIK (BCL2-Interacting Killer, tumor suppressor, apoptosis regulator), FANCE (Fanconi anemia complementation group E), and ECT2 (Epithelial Cell Transforming 2, oncogene). Similarly, unique genes detected under the T2T alignment included several well-studied cancer-related genes, including FOXA1 (key transcription factor in hormone-responsive cancers), BTG2 (tumor suppressor, p53 target, regulator of cell cycle arrest), FOSL2 (AP-1 transcription factor family, proliferation and cancer progression), MAF (oncogenic transcription factor in multiple myeloma), and E2F2 (cell cycle regulator). Similar to RNA-seq, these results suggest that the T2T-CHM13 alignment strategy may improve biological signal, and that alignment with “sponge” sequences may also enhance biologically relevant signal.

We also investigated the effect of “sponge” sequences on single nucleotide polymorphism (SNP) and InDel calls from whole-genome sequencing (WGS) data. We found that data aligned to the hg38 full plus sponge assembly contained only 0.06% SNPs (2,067 out of 3,376,120) and 0.36% InDels (2,582 out of 721,195) without dbSNP build 155 rsIDs (Supplementary Figure S14A, Additional File 24). By contrast, data aligned to the hg38 autosomal assembly contained 0.63% SNPs and 0.92% InDels without rsIDs. Data aligned to the T2T-CHM13 genome assembly had the largest proportion of unannotated SNPs and InDels (6.71% and 52.94%), which is expected given the liftover-based nature of dbSNP155 annotations to T2T. Of the annotated SNPs, only 0.35% were unique to the hg38 full plus sponge assembly, while 5.59% were unique to the hg38 autosomal assembly. The T2T-CHM13–aligned data contained the largest proportion of unique annotated SNPs (23.02%). For InDels, these values were 0.39%, 3.80%, and 10.16% for the hg38 full plus sponge, hg38 autosomal, and T2T-CHM13 alignments, respectively (Supplementary Figure S14B, Additional File 24). We hypothesized that most assembly-specific variants would be false positives. To test this, we examined the proportion of unique SNPs and InDels that overlapped benchmark variants from the Genome in a Bottle (GIAB) consortium. The majority of SNPs were not annotated (98.44% for hg38 full, 89.96% for hg38 autosomal, 99.36% for T2T-CHM13). Similarly, the majority of InDels were not annotated (99.21%, 89.17%, 99.44%; Supplementary Figure S14C, Additional File 24). In summary, these results suggest that alignment to the hg38 genome assembly with “sponge” sequences results in fewer false positives and better identification of annotated SNPs and InDels.

## Discussion

The original publication describing the Blacklist software provides only limited methodological details, which hinders the reproducibility of its results. For instance, key algorithmic parameters such as k-mer size, “binSize,” “binOverlap,” and the single-versus multi-read configuration of Umap mappability files are not described or benchmarked. Additionally, the correlation analysis demonstrating reduced artificial correlations among ChIP-seq data lacks transparency regarding data sources and correlation methods, prompting us to implement our own version of this analysis. Moreover, the data properties influencing exclusion set generation remain unexplored, leaving room for variability across different datasets and software configurations. In this study, we addressed these gaps by reverse-engineering the Blacklist software, fixing an annotation error, providing corrected C++ and Python code, and systematically investigating the properties of various exclusion sets and algorithmic choices. Despite our efforts, we were unable to fully reproduce the exclusion sets provided in the Blacklist GitHub repository.

While we thoroughly benchmarked the properties of exclusion sets, the Blacklist software contains numerous hidden parameters that may influence results. Comprehensive evaluation of these parameters would require advanced parameter optimization methods, such as grid search algorithms [40]. Data preprocessing steps, such as adapter trimming, also warrant investigation. Although we assumed the BAM-derived FASTQ files were pre-trimmed, testing raw FASTQ files could reveal the impact of trimming software. Similarly, peak caller choice is critical in defining high-signal regions. We used MACS3 [20], the latest iteration of MACS2, but alternatives such as Genrich and deep learning-based tools (e.g., LanceOtron [41]) could be evaluated to determine their influence on results.

Exhaustive testing of exclusion set generation tools is beyond the scope of this work. Greenscreen software, despite being Docker-wrapped, was technically challenging to run and featured numerous parameters affecting its pipeline. The algorithm underlying the GreyListChIP R package is unpublished, with internal parameters comparable to those in Blacklist software. While GreyListChIP performed well in reducing artificial correlation among ChIP-seq peaks, we observed some unexplained results, such as missing calls on specific chromosomes. Benchmarking GreyListChIP remains an area for future exploration.

We recommend caution when selecting exclusion sets. While our “High Signal” and “Low Mappability” regions are potential candidates for exclusion, they differ significantly from Blacklist-generated lists and overlap many protein-coding genes and cancer drivers. The “Nordin CUT&RUN” and GreyListChIP-generated lists appear effective at reducing artificial correlations but also overlap cancer driver genes. We provide GreyListChIP-generated exclusion sets for hg38, T2T, mm10, and mm39, together with other exclusion sets generated in this work, in our excluderanges R package [12] but advise applying GreyListChIP to experiment-specific input data.

Using “sponge” sequences at the alignment step appears to be a viable alternative to exclusion sets. We demonstrate that including “sponge” sequences at the alignment step reduces artifact signals comparable to Blacklist-generated lists. The fact that some reads overlapping excludable regions remain aligned even in the presence of “sponge” sequences (Supplementary Figure S12A, Additional File 19) suggests that removing all reads from excludable regions may lead to loss of biologically relevant signal. “Sponge” sequences avoid the problem of poor reproducibility of exclusion sets and potential loss of biological signal by targeting artifact reads at the alignment step. We observed their effectiveness in RNA-seq alignment (negligible effect on gene expression), ATAC-seq (improving biological signal around clinically relevant genes), and in whole genome sequencing settings (minimal loss of genomic variants that are likely false positives), suggesting that including “sponge” sequences as part of genome references may improve biological signal from genomic data generated by any short-read technology.

Technological advancements may reduce the need for exclusion sets and/or “sponge” sequences. Longer sequencing reads (e.g., PacBio, Oxford Nanopore, and the current Illumina 300 bp read length) have a greater likelihood of correct alignment, as we observed (Supplementary Figure S12, Additional File 19). Improved genome assemblies, such as the T2T-CHM13 human reference, also have the potential to enhance biological signal, as demonstrated with experimental omics data (Supplementary Figure S13, Additional File 21). While the hg38 genome assembly remains a gold standard in genomics analyses, we recommend incorporating “sponge” sequences into the assembly to mitigate artifact signals.

Although highly promising, using “sponge” sequences remains a method that is less broadly applicable. This is due to limited understanding of sequences that constitute “sponge” or “decoy” sequences. While these sequences have been assembled for the hg38 human genome [1], they haven’t been created for genome assemblies of other model organisms. Moreover, with the development of the telomere-to-telomere versions of human [16] and mouse [42] genomes these “sponge” sequences may not be necessary. The development of long-read sequencing (Oxford Nanopore Technologies, Pacific Biosciences) may also alleviate spurious alignments, as we have shown when comparing the 36bp and 101bp alignments (Supplementary Figure S12A, Additional File 19). Furthermore, the growing amount of whole genome data and the development of pangenome graph assembly methods has been shown to improve genomic variant discovery, RNA-seq and chromatin immunoprecipitation read mapping [43]. It represents a promising way to alleviate alignment artifacts; however, methods for using graph assembly are still relatively less widely adopted than those using linear genome assemblies. Our future work includes defining “sponge” sequences for other genomes and model organisms as well as exploring the use of the pangenome graph assembly for improving biological signal and eliminate the need for exclusion sets.

## Methods

### Data sources

The original Blacklist exclusion lists for human, mouse, and other species were obtained from the Blacklist GitHub repository (https://github.com/Boyle-Lab/Blacklist/tree/master/lists), using version 2 of the lists. The Kundaje Unified list was retrieved from the ENCODE Project (accession number ENCFF356LFX). The Nordin CUT&RUN exclusion set was originally obtained from Additional File 2 of the corresponding publication [33] and sourced from the excluderanges R package [12]. Unless otherwise indicated, we used lists based on the hg38 human genome assembly.

To replicate the Blacklist exclusion lists, accession numbers of the corresponding BAM files were obtained using metadata provided in the Blacklist GitHub repository (https://github.com/Boyle-Lab/Blacklist/tree/master/lists/metadata) and downloaded from ENCODE. The “Umap” mappability files were obtained from the Hoffman Lab Umap/Bismap Project page (https://hoffmanlab.org/proj/bismap/). Accession numbers of the FASTQ files used to generate these BAM files were identified via the [**PG?**] tag in the header lines of each BAM file and downloaded from ENCODE.

The Gm12878 transcription factor FASTQ files used for correlation analysis were retrieved from ENCODE using the following metadata link: https://www.encodeproject.org/metadata/?type=Experiment&cart=%2Fcarts%2F4b50a1ed-d002-4c76-8401-5ac42d3f2228%2F&files.output_category=raw+data.

Experimental RNA-seq data [44,45] were obtained from GEO accession GSE235167. Experimental ATAC-seq data [46] were obtained from GSE266134.

The whole-genome sequencing (WGS) FASTQ files (ERR194147 for individual NA12878) were downloaded from https://www.ebi.ac.uk/ena/browser/view/ERR194147. The GIAB benchmark SNP/InDel dataset (HG001_GRCh38_1_22_v4.2.1) was downloaded from https://ftp-trace.ncbi.nlm.nih.gov/ReferenceSamples/giab/release/NA12878_HG001/latest/GRCh38/. For hg38 base recalibration, dbSNP build 155 was used in addition to known indels from UCSC (https://storage.googleapis.com/genomics-public-data/resources/broad/hg38/v0/Homo_sapiens_assembly38.known_indels.vcf.gz). For CHM13v2.0, we used the same dbSNP build 155 lifted over by the T2T consortium (https://s3-us-west-2.amazonaws.com/human-pangenomics/T2T/CHM13/assemblies/annotation/liftover/dbSNP.htm) to the appropriate coordinates. All data were accessed on August 12, 2025.

### Generated exclusion sets

Blacklist exclusion sets for the hg38 and mm10 genome assemblies were generated using the BAM files listed in the metadata on GitHub, relevant Umap mappability files, and the original Blacklist software.

To define High Signal regions for the hg38 genome assembly, we generated a signal track by merging 274 paired-end 100/101 bp BAM files (aligned with bwa 0.7.10 sampe) via samtools merge (v1.2). We called local peaks using: `macs3 callpeak --treatment ${in_file} --name 101_local --outdir ${out_dir} --gsize hs --slocal 10000 --llocal 100000 --keep-dup all` (v3.0.1) and selected those with a fold change greater than 99.0% of the fold change range. Regions within 1,000 bp were merged, and regions smaller than 1,000 bp were discarded. This strict filtering ensured the selection of highly significant and long high-signal regions. After generating our High Signal regions, we merged these regions with centromeres from the UCSC Genome Browser, available via the excluderanges R package [12].

To define Low Mappability regions for the hg38 genome assembly, we downloaded the kmer-100 multi-read mappability bedGraph file from the Hoffman Lab website (https://bismap.hoffmanlab.org), which contains base-level mappability values ranging from 0 to 1 (with 1 being highly mappable). We first defined the mappable universe by selecting regions with mappability greater than 0.01. Using bedtools (v2.31.1), regions within 1,000 bp were merged. Inverting the mappable universe resulted in a set of regions with mappability ≤0.01. These regions were then merged within 1,000 bp, and regions smaller than 1,000 bp were discarded. This strategy preserved mappable regions interspersed with short unmappable segments while conservatively selecting long low-mappability regions.

The GreyListChIP R package generates a BAM file-specific exclusion set. We generated GreyListChIP lists for humans by combining 381 36 bp single-end FASTQ files used in the original Blacklist publication, realigning them with STAR to the hg38 genome assembly, and setting the maxGap parameter to 1,000. This list was used in our benchmarks. Additionally, we combined 274 paired-end 100 bp and 101 bp FASTQ files and generated lists for the hg38 and T2T genome assemblies. Similarly, for mice, we used 106 36 bp single-end FASTQ files and 176 50 bp single-end FASTQ files to generate lists for the mm10 and mm39 genome assemblies. All GreyListChIP lists have been added to our excluderanges R package [12].

### Comparison of exclusion sets

We compare two exclusion sets *A* ={*a*_1_, *a*_2_, …, *a*_*i*_} and *B* = {*b*_1_, *b*_2_, …, *b*_*j*_} using the Jaccard count overlap, which is defined as the ratio between the average number of regions in *A* and *B* that overlap with any region in the other set, and the total number of unique regions in the union of *A* and *B*. In mathematical terms, this is given by

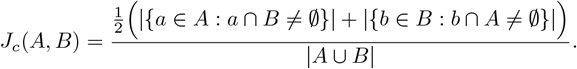

Similarly, Forbes width overlap formula:

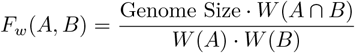

Where *W* (*A*) and *W* (*B*) are the total widths in bases of sets *A* and *B*, respectively, *W* (*A* ∩ *B*) is the total width of the intersection of *A* and *B*, and the “Genome Size” is the total reference genome size (e.g., 2,875,001,522 base pairs for hg38).

Similarity among exclusion sets was visualized using Classical Multidimensional Scaling (MDS) by converting the Jaccard count overlap and Forbes width overlap matrices into distance matrices and plotting exclusion sets using the first two principal coordinates. These distance matrices were also used for hierarchical clustering with the Ward clustering method. All other comparisons (e.g., number and width of overlapping regions) were performed using the GenomicRanges (v1.56.1) R package [47], R (v4.4.0), and Bioconductor (v3.19) [48].

### Aligner test

The original hg38 BAM files were aligned using version 0.7.10 of the bwa sampe (paired-end) and bwa samse (single-end) aligners [49]. In our comparison of exclusion sets generated from BAMs aligned with different aligners, we used the following: bwa-mem2 2.2.1 [49]: `bwa-mem2 mem references/GCA_000001405.15_GRCh38_no_alt_analysis_set.fna.gz <R1.fastq.gz> <R2.fastq.gz> | samtools view -bS - | samtools sort -o <bwamem.bam> -`; bowtie2 2.5.4 [50]: `bowtie2 -x references/GCA_000001405.15_GRCh38_no_alt_analysis_set -U <R1.fastq.gz> <R2.fastq.gz> --local | samtools view -bS - | samtools sort -o <bowtie2.bam> -`; STAR 2.7.11b [51]: `STAR --genomeDir <references/star> --readFilesIn <(bgzip -cd <R1.fastq.gz>) <(bgzip -cd <R2.fastq.gz>) --outSAMtype BAM SortedByCoordinate --alignEndsType Local`, as well as htslib and samtools 1.20 [52].

### Correlation analysis

FASTQ files for each transcription factor were merged, and the resulting files were aligned using STAR 2.7.11b (–alignEndsType Local) to GRCh38_no_alt_analysis_set_GCA_000001405.15.fasta (ENCODE ID: ENCSR425FOI) and to the same reference concatenated with “sponge” sequences [1].

Read count bins for each exclusion set or sponge were computed using deepTools v3.5.6 [31] with the multi-BamSummary bins command (default settings). The Pearson correlation matrix was then obtained using plotCorrelation (–corMethod pearson), and each matrix was clustered and visualized in R using Euclidean distance and complete linkage.

### RNA-seq analysis

We realigned our RNA-seq data [44,45] (GSE235167, n = 144) to two versions of the hg38 human genome assembly from the UCSC Genome Browser. One version included only autosomal, sex, and mitochondrial chromosomes (referred to as “autosomal”), while the other also contained additional random and unplaced contigs, as well as “sponge” sequences [1] (referred to as “full plus sponge”). Additionally, we utilized the T2T-CHM13v2.0 assembly [16] and the corresponding UCSC GENCODEv35 CAT/Liftoff v2 gene annotations downloaded from https://github.com/marbl/CHM13. The data were processed using the Nextflow rnaseq v3.19.0 pipeline [53] with the STAR-RSEM quantification strategy.

### ATAC-seq analysis

The ATAC-seq data from synovial sarcoma SYO-1 cell line (GSE266134) were processed using the nf-core/atacseq v.2.1.2. We used the same genome assemblies as for RNA-seq (hg38 autosomal, hg38 full plus sponge, T2T). We defined peaks differentially accessible under treatment with the small molecule SAE1/2 inhibitor, TAK-981 (subasumstat) vs. untreated cells using macs2 v.2.2.9.1 with default settings for the human genome organism. Peaks were annotated with the annotatePeak function from the ChIPseeker v.1.44.0 R package [54], KEGG pathway enrichment analysis was performed with the enrichKEGG function from the clusterProfiler v.4.16.0 R package [55].

### WGS analysis

Whole-genome sequencing (WGS) data were processed using the nf-core/sarek v3.5.0 pipeline with the BWA-MEM2 setting. We used the same genome assemblies as for RNA-seq (hg38 autosomal, hg38 full plus sponge, T2T). Variant calling was performed using GATK HaplotypeCaller within the nf-core pipeline. Variants (including the GIAB benchmark set) were normalized using BCFtools v1.19 [52], removing duplicates and standardizing variant representation. Variants were annotated with dbSNP 155, and split by type (SNPs or INDELs) using BCFtools. Overlapping dbSNP ids were then used to analyze overlapping variants across assemblies.

### Functional enrichment analysis

Hypergeometric enrichment analysis of transcripts with a non-zero sum of exon widths covered by exclusion regions was performed using the enrichR 3.2 R package [56] and the “KEGG_2019_Human” signature database. Analysis of cancer drivers affected by exclusion sets was conducted using the oncoEnrichR 1.5.2 R package [38], considering the “Cancer associations” analysis.

### excluderanges R package update

We added the GreyListChIP-generated sets to our excluderanges R package [12], making them available within the R ecosystem. We used STAR-aligned sequencing data with 36 bp and 101 bp read lengths for the human genome and 36 bp and 50 bp read lengths for the mouse genome, applying a maxGap = 1,000 merge setting. Specifically, lists for the hg38 and T2T human genome assemblies and the mm10 and mm39 mouse genome assemblies were added.

## Code availability

The modified Blacklist code and scripts to reproduce our analyses and figures are available at https://github.com/dozmorovlab/excluderanges_supplementary. The GreyListChIP data generated in this work is available as a part of the excluderanges R package.

## Funding

This work was supported in part by the George and Lavinia Blick Research Scholarship to MD, the NIH/NCI (R01CA246182, R21CA273779, U54CA283762) grants to JCH, the NIH R56 MH107879 JLM, the DoD grant (HT9425-23-1-1017) and the NCI grant (1R01CA272710-01A1) to ACF.

MGD, JDO, and BPGW wrote the main manuscript text. BPGW, JDO, MN, and MGD performed the experiments and figure preparation. JCH, ACF, JLM provided experimental data and helped design experiments. BPGW, JDO, ALO and KVF analyzed experimental data. MN updated the excluderanges R package. All authors reviewed the manuscript.

